# Third Generation Cytogenetic Analysis (TGCA): diagnostic application of long-read sequencing

**DOI:** 10.1101/2021.08.13.456226

**Authors:** Pamela Magini, Alessandra Mingrino, Barbara Gega, Gianluca Mattei, Roberto Semeraro, Davide Bolognini, Patrizia Mongelli, Laura Desiderio, Maria Carla Pittalis, Tommaso Pippucci, Alberto Magi

## Abstract

Unbalanced Structural Variants (uSVs) play important roles in the pathogenesis of several genetic syndromes. Traditional and molecular karyotyping are considered the first-tier diagnostic tests to detect macroscopic and cryptic deletions/duplications. However, their time-consuming and laborious experimental protocols protract diagnostic times from three to fifteen days. Long read sequencing approaches, such as Oxford Nanopore Technologies (ONT), have the ability to reduce time to results for the detection of uSVs with the same resolution of current state-of-the-art diagnostic tests.

Here we compared ONT to molecular karyotyping for the detection of pathogenic uSVs of 7 patients with previously diagnosed causative CNVs of different sizes and allelic fractions. Larger chromosomal anomalies included trisomy 21 and mosaic tetrasomy 12p. Among smaller CNVs we tested two reciprocal genomic imbalances in 7q11.23 (1.367 Mb), a 170 kb deletion encompassing NRXN1 and mosaic 6q27 (1.231 Mb) and 2q23.1 (408 kb) deletions. DNA libraries were prepared following ONT standard protocols and sequenced on the GridION device for 48 h. Data generated during runs were analysed in online mode, using NanoGLADIATOR.

We were capable to identify all pathogenic CNVs with detection time inversely proportional to size and allelic fraction. Aneuploidies were called after only 30 minutes of sequencing, while 30 hours were needed to call CNVs < 500 kb also in mosaic state (44%). These results demonstrate the clinical utility of our approach that allows the molecular diagnosis of genomic disorders within a 30 minutes to 30 hours time-frame.

## Introduction

Unbalanced Structural Variants (uSVs) are a primary cause of many rare and common human diseases, from Mendelian syndromes to complex diseases and malignancies. uSVs range from aneuploidies to sub-microscopic Copy Number Variants (CNVs) and their detection is a key factor for the diagnosis and the appropriate clinical management of such conditions (Gardner *et al.*, 2011).

Conventional CytoGenetics (CCG), Chromosomal Microarray Analysis (CMA) and Second-Generation Sequencing (SGS) have been vastly adopted to detect uSVs in the clinical settings (Silva *et al.*, 2019; Miller *et al.*, 2010; Pfundt *et al.*, 2017), but each of these approaches has technical limitations impacting significantly on turnaround time and diagnostic yield. First, all are based on time-consuming and laborious experimental protocols, delaying time-to-diagnosis by a two- to a three-weeks period. Second, CCG has a resolution limit of 5-10 Mb, preventing the identification of submicroscopic CNVs. CMA and SGS have much higher resolution (thousands and hundreds of base-pairs, respectively) but their sensitivity is lower for mosaic events. Moreover, SGS strategies are based on the sequencing of short reads (usually 100-150 base-pairs): their resolution strongly depends on the depth of sequencing coverage and they are subject to strong GC-content/mappability biases that hamper accurate identification and correct mapping of CNVs. Consequently, the diagnosis of chromosomal disorders mediated by CNVs is still achieved by the sequential application of multiple techniques, with additional waste of time and costs.

The past few years have seen the emergence of the third-generation sequencing technologies that are characterized by the capacity to generate reads even longer than 10 kb. They show elevated mapping accuracy, providing unprecedented capacity to resolve highly homologous as well as repetitive genomic regions and thus improving CNV detection and mapping. These approaches include Oxford Nanopore Technologies (ONT), that have been already demonstrated to accurately identify CNVs at low depth of coverage (Norris *et al.*, 2016). Moreover, in addition to the ease of use and to the fast sequencing runs, taking from 24 to 48 hours to complete the sequence of a genome, the continuous generation of reads by ONT allows data analysis in real-time, while the sequencing process is ongoing, drastically reducing detection times (Euskirchen *et al.*, 2017).

Here, we describe the application of ONT sequencing to the real-time detection of constitutional or mosaic aneuploidies and recurrent submicroscopic CNVs (microduplications/microdeletions) responsible for known genomic syndromes. First, we obtained sample-to-results time in the 30 minutes - 9 hours range, with improved resolution for mosaic events compared to state-of-the-art techniques (aCGH). In addition, we evaluated sensitivity and specificity of low-coverage ONT sequencing for the identification of pathogenic small CNVs (< 500 kb).

## MATERIALS AND METHODS

### Structural variants

We tested ONT effectiveness for the diagnosis of constitutional genomic disorders on 7 patients with aneuploidies or CNVs of different sizes and mosaic levels, previously identified by conventional cytogenetics or array-CGH (Supplementary Table 1). Larger chromosomal aberrations included trisomy 21 and mosaic tetrasomy 12p. Among small CNVs we tested two reciprocal genomic imbalances in 7q11.23 (1.367 Mb), a 170 kb deletion encompassing NRXN1 and mosaic 6q27 (1.231 Mb) and 2q23.1 (408 kb) deletions. DNA samples were extracted from blood specimens by the QIAamp DNA Blood Mini Kit (QIAGEN). Concentration and purity were checked through the Qubit fluorometer and the Nanodrop spectrophotometer (ThermoFisher Scientific), respectively.

### aCGH

Submicroscopic CNVs were identified in patients referred for genetic diagnosis through the 8×60k ISCA platform (Agilent Technologies), with a mean actual resolution of about 120 kb. The same aCGH design was then used to define genomic breakpoints and estimate cellular fractions of trisomy 21 and tetrasomy 12p. The standard protocol indicated by manufacturer has been applied. aCGH data have been analyzed by the Cytogenomics 5.0 software (Agilent Technologies) and aberrations were called by the ADM1 algorithm with threshold set to 6.0. Cellular fraction of genomic alterations was estimated through the formula suggested by (Valli *et al.*, 2011), with average log2 ratio provided by ADM1 calculation.

### Nanopore sequencing

The sequencing library preparation was performed according to the Oxford Nanopore Technologies (ONT) manufacturer’s protocol for genomic DNA (Supplementary Methods). The seven Nanopore libraries were sequenced on a GridION 5X device for 48 hours to maximize the total throughput, using the latest ONT flow cell chemistry. The wet and dry phases of the experimental protocol are illustrated in Supplemental Figure 1. Metrichor software was used for base calling.

Unbalanced chromosomal alterations and CNVs were detected with Nano-GLADIATOR tool (Magi *et al.*, 2019) (https://sourceforge.net/projects/nanogladiator/). Nano-GLADIATOR is a software tool that allows to analyze WGS ONT data to perform CNVs detection and allelic fraction prediction during the sequencing run (“On-line” mode) and once the experiment is completed (“Off-line” mode). NanoGLADIATOR was used in on-line mode at several time points (from 30 min to 48 h as shown in Supplementary Fig. 1) to detect alterations during the sequencing process and in off-line mode to evaluate the resolution of complete ONT runs for genomic alterations of different sizes. Detection threshold was set to 2 consecutive windows. At the end of each sequencing run, raw data files were analyzed with PyPore package (Semeraro *et al.*, 2019) to check sequencing run quality.

The number and the size of false positive (FP) CNVs were evaluated at different analysis window sizes and at each time point. FP obtained at the end of sequencing (48 h) with the 50 kb analysis window were evaluated further: we excluded FP events with size and cellular fractions under the detection thresholds of the aCGH platform used (120 kb and 20%, respectively) or mapping in genomic regions not covered by the aCGH design. In addition, we filtered out also deletions and duplications entirely located within known polymorphic regions (indicated as CNVR within the Database of Genomic Variants, DGV, http://dgv.tcag.ca/dgv/app/home) and/or overlapping segmental duplications, impairing hybridization.

### Quantitative PCR (qPCR)

qPCR was performed through the Universal Probe Library (UPL) system (Roche, Basel, Switzerland). Two primer pairs (sequences available on request) were designed (Assay Design Center, http://lifescience.roche.com/) to amplify two genomic regions mapping within the CNV. The DNA of two healthy individuals was processed along with patient’s DNA. Reactions were performed in triplicate for each primer pair, simultaneously amplifying the RPPH1 gene as disomic reference (TaqMan Copy Number Reference Assay; Life Technologies, Grand Island, NY). The ΔΔCt method was applied to real-time data to obtain a relative quantification of copy number of analyzed genomic regions.

## RESULTS

For each sample, we completed the process from DNA extraction to flow cell load in about 3 hours (Supplementary Fig. 1). All known pathogenic imbalances were detected by ONT at different times depending on their size. After starting the sequencing process, Nano-GLADIATOR was used in on-line mode by setting time points for data analysis as shown in Supplementary Fig. 1. With sequencing data generated at the first time point (< 100 k reads at 30 minutes, Figure. 1.a), we were capable to detect aneuploidies and large chromosomal alterations (Figure 1.b).

**Figure 1:**
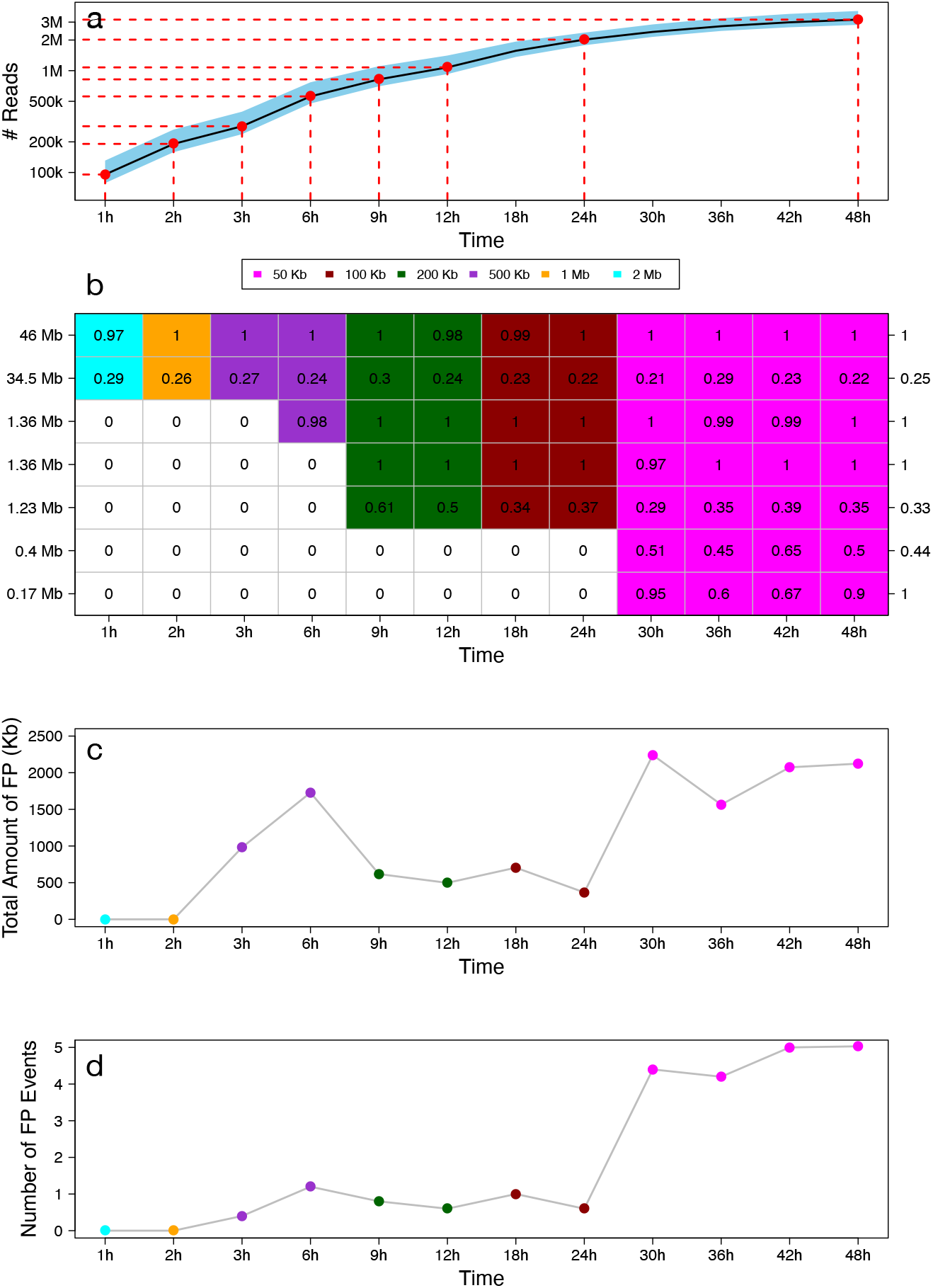
Nanopore sequencing throughput and CNVs detection accuracy. Panel (a) shows the average number of reads (both pass and fail reads) generated during a typical MinION/GridION sequencing run with R9.4 flow-cells. Data are averaged across the seven runs performed for this work with R9.4.1 flow-cells on GridION device. Panel (b) reports the capability of ONT data and NanoGLADIATOR to detect genomic alterations of different size and predict their cellular fraction as a function of sequencing time. In x axis is reported the sequencing time, while in y axes are reported the genomic alteration characteristics (Left y-axes the size of the alteration, right y-axes the cellular fraction predicted by aCGH). Colors of the heatmap indicate the window size used to detect the genomic alterations, while the number in each cell indicates the predicted cellular fraction. White cells indicate that NanoGLADIATOR was not capable to identify the alteration. From bottom to top are reported samples with pathogenic CNV of increased size: 23091, 24327, tp_1406, 23023, 20686, 25488 and 23984 (see Supplementary Table 1). Panels (c-d) reports the average size (c) and number (d) of false positive calls made by NanoGLADIATOR at different time points and with different window size. Each point of the plot is averaged across the seven runs performed for this work.

Resolution improved proportionally to the duration of the sequencing process and to the parallel coverage increase: microdeletions/microduplications of about 1 Mb were detected in 6-9 hours (500 k-1 M reads), while those smaller than 500 kb were visible after 30 hours, (Figure 1.b). Remarkably, we found our approach to have higher sensitivity for mosaic CNVs compared to aCGH. In a patient (25488) presenting with Pallister-Killian Syndrome (PKS) we were able to identify a chromosome 12p amplification having an estimated cellular fraction of 23%, that was missed by aCGH aberration calling algorithms (ADM1 and ADM2) but recognized after a visual inspection of the aCGH profile.

To further study ONT sensitivity and specificity, we then compared all ONT calls generated after 48 hours of sequencing, with a final average coverage of 5-6X, to those provided by aCGH in each patient in addition to the pathogenic ones. We evaluated genomic regions, size and cellular fractions of discrepant calls. We did not identify false negative calls, namely CNVs detected exclusively by aCGH. CNVs with no overlap with aCGH findings were considered false positive events. The average number of FP correlated with sequencing time and the windows size used for analysis detection (Figure 1.c). Within the first 2 hours of sequencing run, with windows size of 1 and 2 Mb, we had evidence of no FP events. From 3 to 24 hours (window size from 500 Kb to 100 Kb), we identified on average one FP event, with a maximum of total size FP reached at 3-6 hours of sequencing with the 500 kb window (Figure 1.d).

From 30 hours, through the application of the 50 kb window, the number of FP events grew up to 4-5 on average, covering up to 2.5 Mb and showed a linear inverse correlation between cellular fraction and size (Supplementary Fig. 2). From all FP detected at the end of the sequencing process we excluded those undetectable by aCGH or mapping in polymorphic regions, obtaining a total of 15 FP calls in 6 samples (Supplementary Table 2). These FPs were all at a mosaic state, generally with estimated low cellular fractions, except one (the homozygous deletion on chromosome 6 in sample 23984) that was confirmed to be a false call as it was not validated by qPCR.

Taken as a whole, these results show that increasing resolution of our approach by narrowing the window size has only limited impact on specificity resulting in a still manageable burden of FP events.

## DISCUSSION

Genetic diagnosis is a fundamental step for the clinical management of patients with genomic disorders and their families, allowing the identification of healthy carriers, a precise definition of recurrence risk, the planning of specific treatments and surveillance screenings when prognosis is known. The greatest benefits are obtained when diagnosis is rapid.

By analyzing genomic DNA from seven patients, we demonstrate that ONT provides fast and accurate detection of uSVs with different sizes and cellular fractions. ONT resolution for uSVs is higher compared to cytogenetic analyses and at least similar to that provided by aCGH, with the additional advantage of real-time detection of aberrations and then rapidity of the whole analysis process. Large chromosomal anomalies, usually detected by karyotyping after several days of sample processing and metaphase analysis, can be identified by ONT with a sequencing run of 30 minutes. Considering about three hours to process the sample from DNA extraction to library preparation, diagnosis of cytogenetic alterations and recurrent microdeletion/microduplication syndromes can be made in one day. CNVs smaller than 500 kb are detected after 30h of sequencing but the turnaround time for their diagnosis, estimated in 2 days, is shorter than that required by aCGH (at least 3 days) with the further advantage of an easier wet-lab procedure. Potentially, ONT resolution and sensitivity outdoes that of aCGH thanks to its nucleotide-level resolution and to more homogeneous genome analysis, sequencing also structurally complex or high GC-rich regions that are not covered by aCGH for hybridization problems.

However, false positive rate should be carefully considered, especially for a perspective diagnostic application of ONT. False positive events concern mostly very small CNVs and potential low-mosaic imbalances, detected at the highest resolution applied (50 kb analysis window). With a focus on cytogenetic aberrations (> 5 – 10 Mb), reliably detected by 1 and 2 Mb analysis windows, no FP appears at any coverage, making the diagnosis of large CNVs very specific, easy and rapid. Analysis windows of 100-200 kb gave the best compromise between specificity and sensitivity, generating very few FP and allowing the detection of recurrent microdeletions/microduplications responsible for known genomic disorders in 9-18 hours. Increasing analysis resolution for the identification of CNVs < 500 kb, the number of FP raises to an average of 5 events per sample, even less if non-clinically relevant CNVs are excluded. Although this rate is still manageable, false calls should be easily discriminated from true deletions/duplications, to assure a straightforward analysis of ONT data in a diagnostic setting. A greater sequencing depth is probably required to stabilize calls from 50 kb analysis windows and detect very small CNVs with high specificity. Nevertheless, since all but one FP events has cellular fraction < 55% in our dataset, we can confidently suppose that low coverage ONT sequencing has good accuracy for the detection of germline CNVs smaller than 500 kb.

We also demonstrated that ONT can identify mosaic CNVs, even of small sizes (1231 and 408 kb, Supplementary Table 1), with a higher sensitivity compared to aCGH, which was unable to call a complete 12p duplication/triplication with a cellular fraction of about 23%. Pathogenic mosaic structural variants have been identified in approximately 0.3-0.77% of patients affected with developmental disorders (Hoang *et al.*, 2011; King *et al.*, 2015). Since the sensitivity of current diagnostic tests for their detection is limited, with a minimum cellular fraction detectable by aCGH of about 25% (Hoang *et al.*, 2011), the frequency of mosaicisms as cause of genetic diseases may be underestimated. Moreover, mosaicism may be missed in parents of children with apparently de novo pathogenic CNVs, leading to an incorrect estimation of recurrence risk and to an improper management of future pregnancies.

Greater sensitivity for mosaic detection will also be useful for the diagnosis of genetic disorders, such as Pallister-Killian syndrome, caused by supernumerary chromosomes which are negatively selected in phytohaemagglutinin (PHA) stimulated peripheral blood lymphocytes, requiring skin biopsy for chromosomal analysis, and are lost over time in actively dividing tissues, such as bone marrow, impacting diagnosis in older children and adults (Izumi *et al.*, 2014). High rapidity, accuracy, resolution and sensitivity for uSV detection, even at mosaic state, make ONT an efficient diagnostic tool that can identify cytogenetic and microscopic anomalies in a single analysis, considerably reducing turnaround times. Perspective studies on large cohort of patients addressed to karyotyping and aCGH are needed to test ONT performance in routine clinical settings.

## Data Availability Statement

Nanopore sequencing data are not publicly available since patients did not consent for genetic data sharing.

## Ethics declarations

### Contributions

Conceptualization: Alberto Magi, Tommaso Pippucci. Investigation: experiments: Pamela Magini, Alessandra Mingrino, Patrizia Mongelli, Laura Desiderio, Maria Carla Pittalis. ; computational analyses: Barbara Gega, Pamela Magini, Roberto Semeraro, Gianluca Mattei, Davide Bolognini. ; Methodology: Pamela Magini, Alberto Magi, Tommaso Pippucci. Writing—original draft: Pamela Magini, Alberto Magi, Tommaso Pippucci. Writing—review editing: all authors.

### Ethics declaration

Individuals were referred by clinical geneticists for genetic testing as part of routine clinical care. All patients enrolled and/or their legal representative signed informed consent for research use.

