## Supplementary Figures for "Third Generation Cytogenetic Analysis (TGCA): diagnostic application of long-read sequencing"

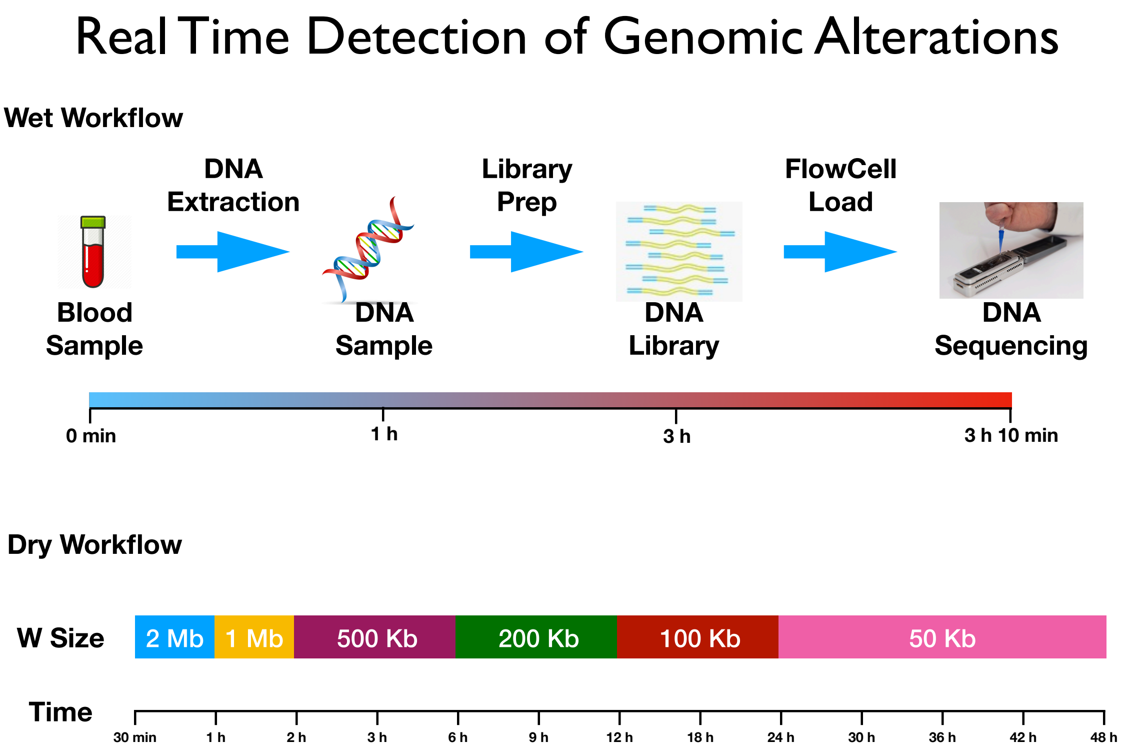


Figure 1: Timeline of the experimental process. In Figure is reported the timeline of the wet and dry workflow. Wet workflow consists of DNA extraction, library preparation and FlowCell loading. Dry workflow consists in running NanoGLADIATOR during the sequencing process by using different window size at different time steps: 2 Mb within the first hour, 1 Mb between 1 h and 2 h, 500 kb between 2 h and 6 h, 200 kb between 6 h and 12 h, 100 kb between 12 hand 24 h and 50 kb between 24 h and the end of the sequencing process.

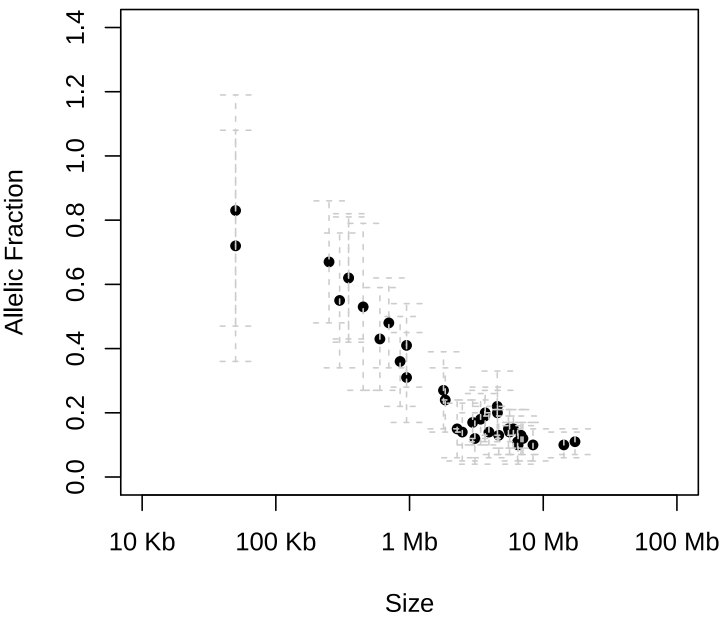


Figure 2: Correlation between size and allelic fraction of FP calls. In figure are reported the allelic fraction and size of FP calls made by NanoGLADIATOR in the analysis with window size 50 kb.
